# Learning to predict based on self- versus externally induced prediction violations: a direct comparison using a Bayesian inference modelling approach

**DOI:** 10.1101/2022.11.15.516578

**Authors:** E.A. Boonstra, H.A. Slagter

## Abstract

Predictive processing is quickly gaining ground as a theory of perception and attention. From this perspective the brain is cast as an organism’s predictive model of how its world works and will continue to work in the future. However, research on the brain’s predictive capacities remains beholden to traditional research practices in which participants are passively shown stimuli without their active involvement. The current study is an investigation into ways in which self-generated predictions may differ from externally induced predictions. Participants completed a volatile spatial attention task under both conditions on different days. We used the Hierarchical Gaussian Filter, an approximate Bayesian inference model, to determine subject-specific parameters of belief-updating and inferred volatility. We found preliminary evidence in support of self-generated predictions incurring a larger reaction time cost when violated compared to predictions induced by sensory cue, which translated to participants’ increased sensitivity to changes in environmental volatility. Our results suggest that internally generated predictions may be afforded more weight, but these results are complicated by session order and duration effects, as well as a lack of statistical power. We discuss the limitations of our study preventing us from replicating previous research, and ways to remedy these shortcomings in future studies.

## Introduction

The study of perception and attention in living organisms is most often conducted from an outside-in perspective, from which primacy is afforded to information and stimuli impinging on sensory epithelia. For example, tasks are used in which external cues tell participants where to attend. This perspective or approach becomes problematic if we recognize that living organisms maintain internal dynamics of their own, based on which they perceive, attend, act, and operate. Indeed, it has been argued that an inside-out perspective, where an organism’s internal states are afforded primacy, is a more appropriate starting point for the study of brain and behavior (Buzsáki, 2019).

An inside-out perspective is inherent to research on self-elicited action-effects. Already in the 1970s, the seminal study by Schafer and Marcus (1973) reported that self-delivered auditory and visual stimuli attenuated the amplitude of event-related potentials (ERPs) by respectively 50% and 28%, compared to when stimuli were machine-delivered. Since then, research on the directional relationship from action to perception has been framed in terms of agency; revolving around the question how and to what degree the system ascribes the cause of sensation to itself, in contrast to unsolicited stimulation from outside (Synofzik et al., 2013; Wen, 2019) and action-effects (Elsner & Hommel, 2001; Hommel, 1996, 2019; Hommel & Elsner, 2009; Prinz, 1997). In the auditory domain in particular, the finding that self-generated tones are accompanied by attenuated early ERPs such as the N1 and P2 has been replicated many times (Korka et al., 2022), but the behavioral effect is less conclusive: perceptual sensitivity has been found to both decrease (Weiss et al., 2011; Weiss & Schütz-Bosbach, 2012) and increase (Reznik et al., 2014; Reznik & Mukamel, 2019). While the auditory domain is more extensively studied, similar neural effects have been observed for the visual domain (Cardoso-Leite et al., 2010; Nittono, 2004; Nittono et al., 2003; Vasser et al., 2019; but see Schwarz et al., 2018). While the exact workings of the relationship between action and perception are unclear as of yet, it is clear that stimuli stemming from self-generated predictions are processed differently from externally induced ones.

An inside-out perspective is also part and parcel of predictive processing approaches to the study of brain and behavior (Clark, 2013; Friston, 2010; Hohwy, 2013). From these perspectives, the brain instantiates an organism’s predictive model of its outside world, which upholds expectations regarding how this world works and how it will continue to work in the future. This model undergoes changes as the organism is confronted with violations to its expectations; violations it tries to minimize overall. This process of model-updating and prediction error minimization can be cast as approximate Bayesian inference (Da Costa et al., 2021; Knill & Pouget, 2004; Pouget et al., 2013). From this perspective as well, the violation of internally and externally generated predictions are thought to be weighted differently (Brown et al., 2013), but to our knowledge no study has made this comparison directly for the visual domain within the context of Bayesian predictive perceptual processing.

We set out to test this difference within the context of the Hierarchical Gaussian Filter (HGF); a computational model implementing approximate Bayesian inference linking three hierarchically ordered levels capturing not only trial-by-trial probability estimates, but also the inferred volatility of an agent’s environment (Mathys et al., 2011; Mathys et al., 2014). We rely heavily on earlier work in which this model was used to show how model-updating changed over time in the context of a spatial attention task (Vossel, Bauer, et al., 2014; Vossel, Mathys, et al., 2014). In these studies, a cue indicated whether a target would appear left or right of fixation. Crucially, the probability with which the cue was valid changed over the course of the experiment across three levels (50, 69, 88%) unbeknownst to participants. It was found that participants were not only faster with eye movement responses on valid compared to invalid (e.g., cue indicating left, target appearing right) trials, but this effect was stronger on blocks where the probability of a valid cue was larger. In terms of modelling, they found an optimal model aligned with the proposal by Feldman and Friston (2010), suggesting that in general the precision of prediction errors drive changes in model-updating, to the point that higher precision results in more rapid updating, and where in the case of this particular model, the trial-by-trial precision estimate of the first-level prediction drives responses. The importance of contrasting this setup with an action condition in which predictions are self-generated comes from the idea that through action, a predictive system can actively test the validity of its internal model. Instead of being at the whim of external stimulation (passively awaiting what comes in), the system can generate sensory information to actively test its models, facilitating the speed at which reliable models can be produced.

Our main research question is as follows: how does the response to induced predictions and their violation differ when these predictions stem from externally generated sensory cues compared to internally generated action?

We first report a pilot study in which we transposed the experimental design by Vossel and colleagues (2014) to an action-effect paradigm. Our pilot was the same as the original study, except that spatial predictions were self-generated by participants’ own actions instead of external cues. We expected to replicate Vossel and colleagues (2014) by finding slower eye movement responses for invalid compared to valid trials, and that this difference would be larger for higher probability blocks. Like Vossel and colleagues (2014), we expected an optimal “precision” response model to account for these findings. An indirect comparison with Vossel et al.’s findings showed that the RT cost for violated predictions were larger when this prediction stemmed from self-generated action, in our pilot study, which shaped our hypotheses for our main within-subject experiment in which participants completed an action and cue version of the same task on different days. This allowed us to critically, directly test if these conditions differ by keeping constant as many confounds as possible.

We expected to replicate and extend our pilot finding; that the violation of predictions through invalid trials would incur a larger reaction time cost when a prediction stems from action rather than from a sensory cue. Moreover, in terms of HGF modelling, we expected to see these effects reflected in a larger belief-updating parameter (ω_2_, see below) for the action condition. This would support the notion that perceptual predictions are afforded more weight when they are self-generated.

## Methods

### Participants

Twenty participants were recruited from the Vrije Universiteit Amsterdam subject pool for the pilot, while thirty-five participants were recruited for the main experiment. After exclusion based on pre-specified criteria (see below), the final sample for the pilot consisted of 19 participants (mean age 21.2, range 18-31 years, 16 women), while for the main experiment 25 remained (mean age 19.8, range 18-23 years, 20 women). All participants reported normal or corrected-to-normal vision, provided written informed consent prior to participation, and received either a monetary reward or course credit for participating. The experiment was approved by the ethics review board of the Faculty of Behavioral and Movement Sciences of the Vrije Universiteit Amsterdam.

### Procedure and experimental paradigm

In the pilot study, participants performed a spatial predictability task, in which the predictability of the sensory outcomes of their own actions (a button press) varied across trials. In the main experiment, participants came to the lab for two sessions, and either performed this task or a task in which the spatial predictability of a visual cue was varied across trials (cf. Vossel et al., 2014). In both tasks, participants had to move their eyes as quickly and as accurately as possible to a target appearing either to the left or right of a central fixation cross. The design and analyses of the main experiment were preregistered (https://doi.org/10.17605/OSF.IO/V6DJ2), unless otherwise noted.

Stimuli were presented on a CRT monitor (1680 × 1050 pixels, 75 Hz). Eye movements were recorded using an EyeLink 1000 Plus eye tracker (SR research, Ontario, Canada). Participants viewed the screen through a chinrest at a distance of 70 cm.

Participants were instructed to fixate centrally on an ABC fixation dot as defined by Thaler and colleagues (2013) (.55° in size) until a target appeared. Once the eye tracker registered fixation, the trial would start 500 ms later by the presentation of two squares on the horizontal midline (1.9° in size, 8.1° from fixation) to the left and right of the fixation dot, indicating to participants either an impending cue or that they were to initiate an action, depending on the condition. In the action condition, participants initiated trials by freely choosing one of two buttons (“s” and “k” on a standard keyboard) with their index fingers. The left (“s”) button most often gave rise to a target on the left, while the right (“k”) button had the same function for a target on the right, exploiting a natural spatial association (Leuthold, 2011; Simon & Rudell, 1967), also present for the left and right arrow head cues used in the cue condition (Figure 1a). Participants were instructed to use both buttons approximately evenly, and to avoid simply alternating between buttons. On average participants took 534 ms (SD: 134 ms) to initiate an action. In the cue condition, a cue was presented after 500 ms for 200 ms without the participant’s involvement. 800 ms after an action or a cue, a target in the form of a gabor patch (1.3° in size) appeared in one of the two placeholders to which participants were instructed to move their gaze as quickly and as accurately as possible. Once participant’s gaze reached the target, or after 1700 ms, the trial would end and the next trial would start (see Figure 1).

**Figure 1.**
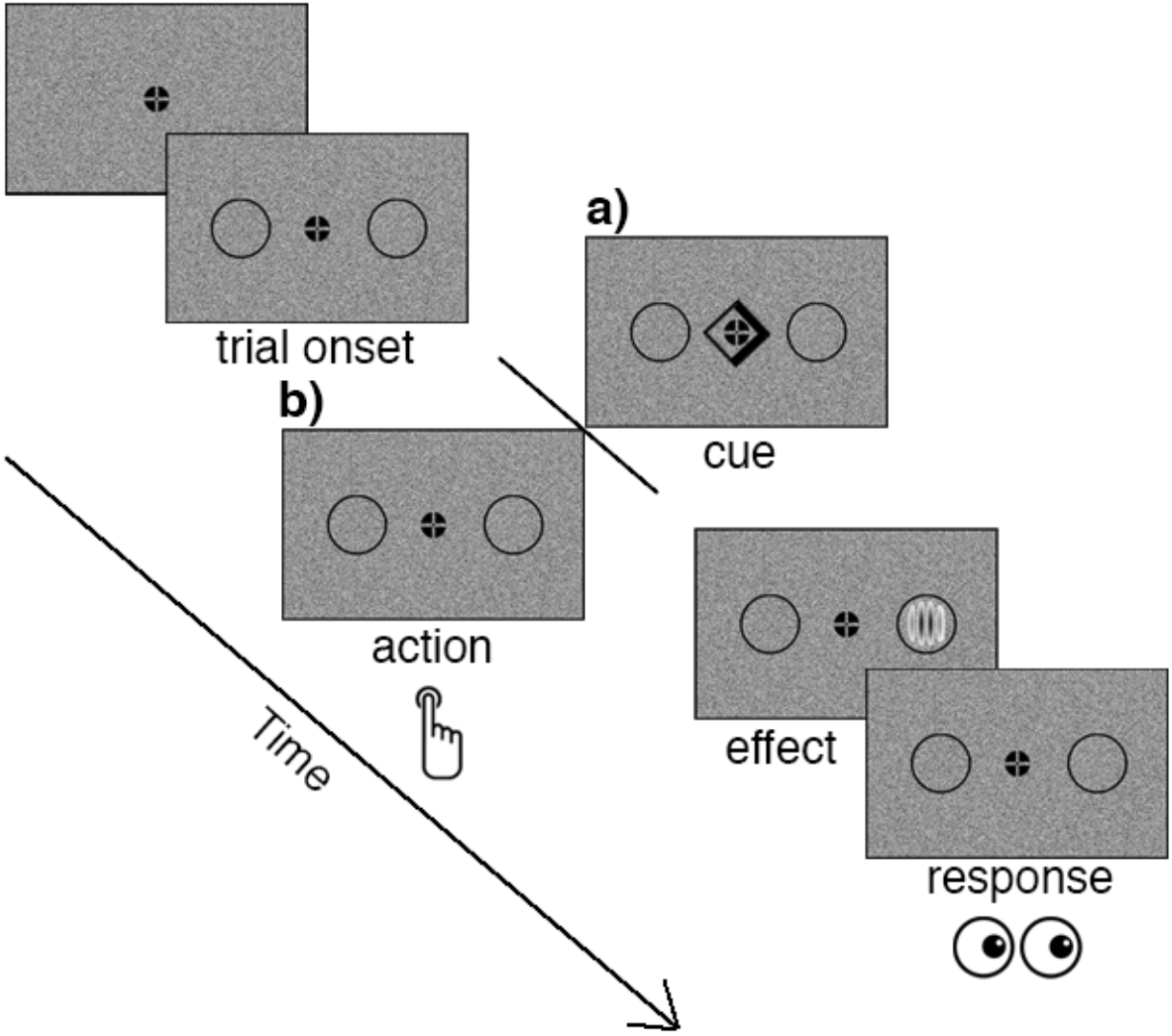
Experimental procedure for one trial in the **a)** cue and **b)** action condition. Depending on the condition, a cue or an action predicted whether a target was likely to appear left or right of fixation or equally often at either location, and participants had to move their gaze to the target when it subsequently appeared.

One session took approximately 1.5 h, in which participants completed a practice block of 36 trials, as well as 612 experimental trials, evenly divided into 18 blocks of either 32 or 36 trials. Trials in the cue condition were interspersed with 108 “null-trials”, where only the baseline display (fixation and squares) were shown to interrupt the flow of trial presentation (cf. Vossel et al., 2014). Crucially, blocks of trials varied unbeknownst to participants in how likely the target was to follow either cue or action. Said differently, when a cue indicated left or when a participant pressed the left button, it was uncertain whether the target would appear on the left side. This validity probability varied block wise at 88, 69, and 50%. The practice block was steady in terms of validity probability at 88%. We used a within-subject design where all participants completed two sessions on different days with at least one day in between. The pilot consisted only of the action session.

### Eye tracking and preprocessing

We used the same analysis pipeline for all the data reported in this paper. All data were analyzed offline. In line with Vossel and colleagues (2014), the start and endpoints of saccades were defined using a velocity-based algorithm, except we used the data-driven method developed by Nyström and Holmqvist (2010) to account for individual differences and thereby maximize sensitivity for saccade detection. We calculated saccade latency (time between target presentation and the start of the first eye movement) and landing position of the first saccade for every trial. In line with Vossel and colleagues (2014), only the first saccade was analyzed.

A target was considered reached when participant’s gaze traversed at least two-thirds of the distance towards the target for at least 10 ms. Trials were excluded when a saccade started out more than 1° from fixation, when saccadic RT was smaller than 90 ms, when less than two-thirds of the distance was traversed to one of either target locations, when no saccade could be detected, when a trial contained more than 20% missing values, and when a response was incorrect. Individual participants were excluded from analyses entirely when we found more than 50% of their data to be missing. In this way we excluded one participant for the pilot and two for the main experiment to arrive at our final samples.

### Data analysis *Pilot experiment*

Our main dependent measure was response speed (RS): the inverse of saccadic reaction time due to its normal distribution (Vossel, Mathys, et al., 2014). To verify that the probability and validity manipulations worked also when predictions were internally generated through action, we ran a 3 (Probability; 88/69/50%) x 2 (Validity; valid/invalid) repeated-measures ANOVA with RS as the dependent measure.

### Main experiment

To test our main question, whether internally generated predictions differ from externally induced ones, we conducted, as preregistered, a 3 (Probability; 88/69/50%) x 2 (Validity; valid/invalid) x 2 (Expectation; action-/cue-induced) repeated-measures ANOVA with RS as the dependent measure. We computed two additional ANOVAs to control for possible confounds. One with the between-subject variable Session Order added, and one in addition to our preregistration, but in line with Vossel and colleagues (2014), with the within-subject variable Time (first/second half of the experiment). We report Greenhouse-Geisser corrected p-values when the assumption of sphericity was violated, as indicated by decimal denoted degrees of freedom.

To evaluate evidence in favor of our (null) hypotheses, we conducted Bayesian statistics. For each reported frequentist ANOVA, we report the Bayes factor corresponding to the inclusion of a factor or interaction within the model in question (shortened to BF_incl_), compared to equivalent models stripped of the effect. For example, BF_incl_ = 10 indicates that a model including the factor in question is ten times more likely given the data compared to a model without the variable. Conversely, BF_incl_ = .1 indicates that a model without said effect is ten times more likely given the data. In addition to frequentist t-tests, we report the Bayes factor corresponding to the relative likelihood of a difference between conditions versus no difference (shortened to BF_10_). All Bayesian statistics were conducted using JASP version 0.16.2 (Love et al., 2019).

Data visualization was performed with the help of raincloud plots (Allen et al., 2019), which include the mean, individual data points, as well as the overall distribution of the measure in question.

### HGF

To formalize subject-specific trial-by-trial belief updating, we employed a Hierarchical Gaussian Filter (HGF) as implemented in the TAPAS toolbox (http://www.translationalneuromodeling.org/tapas/) (Frässle et al., 2021; Mathys et al., 2011; Mathys et al., 2014) for MATLAB (2019a, The MathWorks, Inc., Natick, Massachusetts, United States). The HGF is a Bayesian hierarchical learning model consisting of a perceptual part in which a probability is estimated (in the case of the current study the probability of a valid cue/action), and a response part in which this probability estimate is transformed into RS (see Figure 2). An important advantage of this model is that because it implements approximate (or variational) Bayesian inference, it avoids the computationally costly procedure of Markov chain Monte Carlo (MCMC) sampling, a commonly used Bayesian modelling framework (cf. Wagenmakers & Lee, 2014).

**Figure 2.**
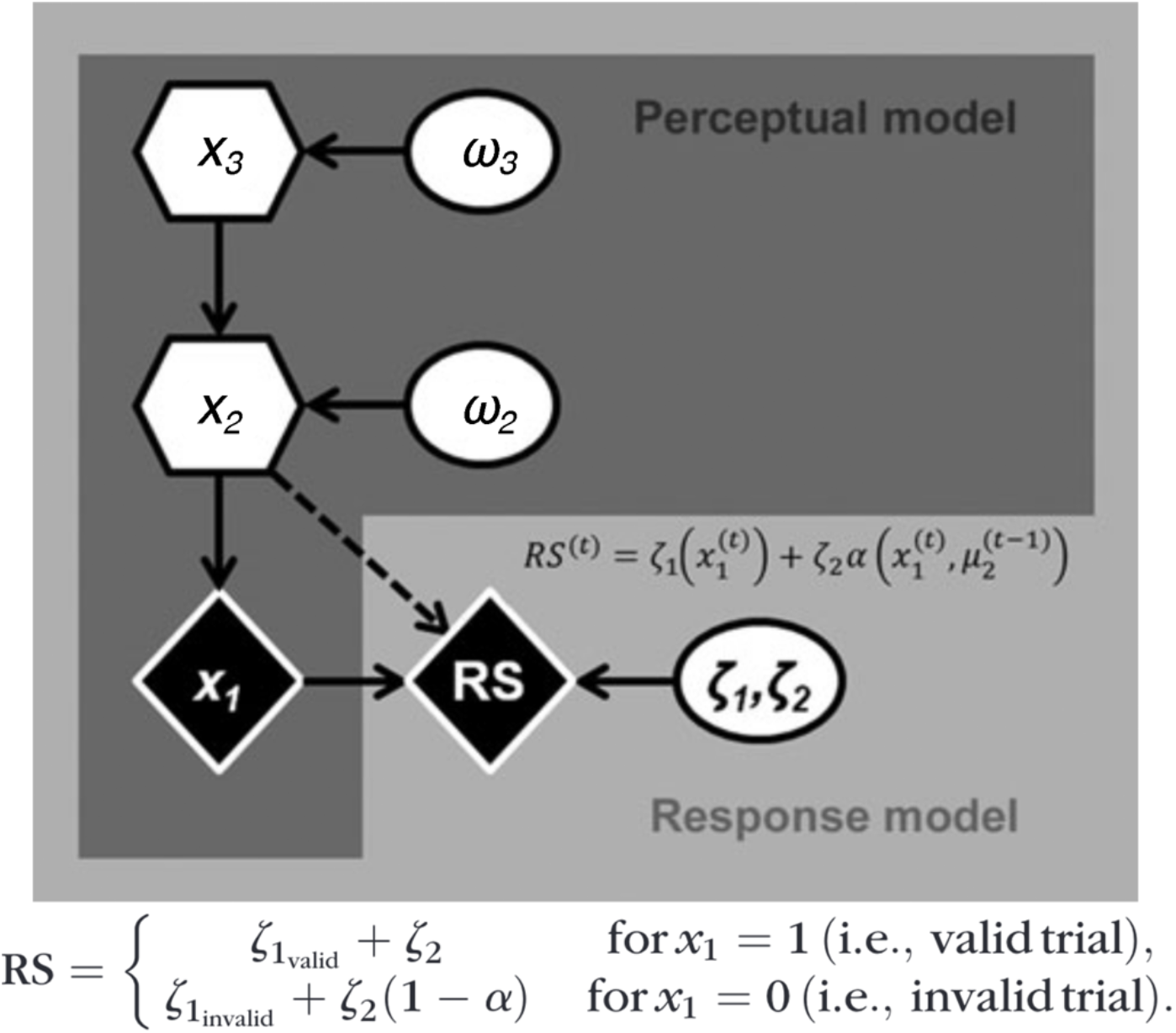
The hierarchical gaussian filter (HGF). Adapted from Vossel and colleagues (2014).

The first level (x_1_^(t)^) of the perceptual model comprises a Bernoulli distribution from which the trial-by-trial probability estimate of a valid cue/action is derived. This level is governed by the level above (x_2_^(t)^). The pace at which this second level changes is governed by the third and final level above (x_3_^(t)^) as well as a subject-specific fixed parameter ω_2_. The step size at which the third level changes is governed by a second subject-specific parameter ω_3_ (denoted as *ϑ* in earlier implementations of the HGF (Mathys et al., 2014; Vossel et al., 2014)). Both the second and third level change as a random Gaussian walk. Said differently, ω_2_ reflects the speed of belief updating about trial-by-trial cue/action validity, while ω_3_ reflects the speed at which the stability of action/cue validity is updated.

Subject-specific beliefs about trial-by-trial cue/action validity and volatility (posterior densities in relation to the hidden states x^(t)^) are inferred from observable behavior (measured RT) by inverting the perceptual model. The sufficient statistics of the subject-specific posterior beliefs are denoted by *μ*^(t)^ (mean), *σ*^(t)^ (variance), and *π*^(t)^ = 1 / *σ*^(t)^ (precision).

The response model captures how an agent’s belief translates to a decision (Daunizeau et al., 2010), or more specifically how the probability estimate at the first level (xι^(t)^) translates to response speed. The response model comprises two parameters ζ_1_ and ζ_2_ which reflect the intercept and slope, respectively, of the linear relationship between the attentional factor α and posterior belief *μ*_1_ (see Figure 2). This attentional factor reflects a proportion of attentional resources allocated to one of two stimulus locations. Three response models were defined by Vossel and colleagues (2014) which differed in how α was computed: 1) a “belief” model in which RS is a linear function of the probability estimate at x_1_^(t)^, 2) a “precision” model where α was determined by a sigmoid transformation (s) of π^(t)^, the precision of the prediction at the first level (Feldman & Friston, 2010; Vossel, Mathys, et al., 2014), and 3) a “surprise” model where α is a nonlinear function of Shannon surprise (Bestmann et al., 2008; Vossel, Mathys, et al., 2014). See Figure 3 for the relationship between α and the probability estimate *μ*_1_ at the first level. Out of these three models, we selected the most likely one given our data through Bayesian model selection (Stephan et al., 2009).

**Figure 3.**
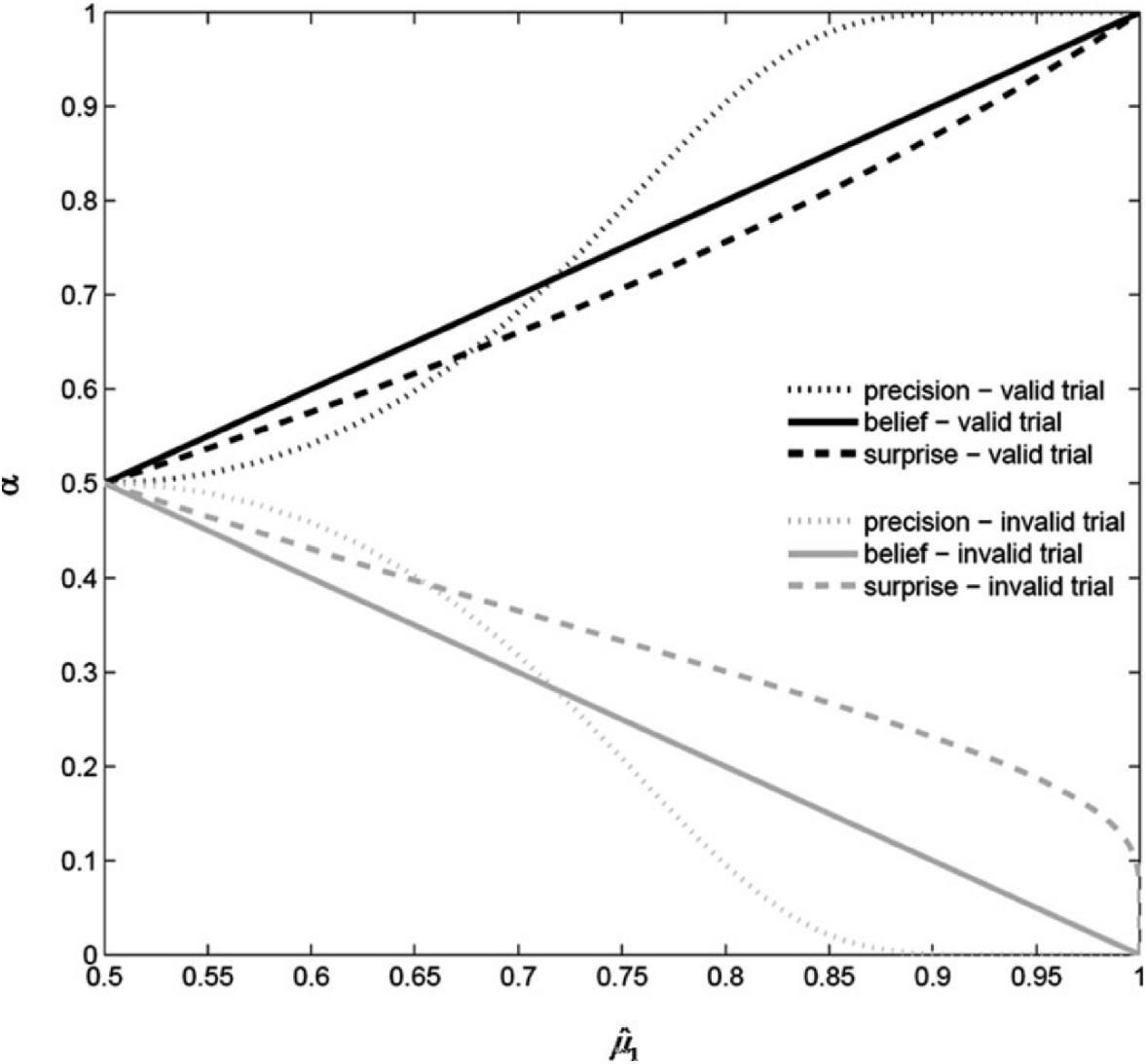
The relationship between attentional factor α and posterior belief *μ*_1_ for all three response models. Adapted from Vossel and colleagues (2014).

## Results

### Pilot

For the pilot, 22.9% of data was excluded from further analyses. Similar to Vossel et al (2014), 19.3% of trials were excluded because the saccade started more than 1° from fixation, 1.1% because saccadic RT was smaller than 90 ms, 1.8% due to less than two-thirds of the distance being traversed to one of either target locations, .09% because no saccade was detected, .4% because trials consisted of more than 20% missing values, and .19% due to incorrect trials.

#### Eye movement behavior

Our pilot study aimed to verify that like increasing external cue validity (Vossel et al., 2014), increasing the predictability of an action outcome would induce a larger RS cost when this prediction was violated. This was confirmed by our pilot data. The 3 (Probability; 88/69/50%) x 2 (Validity; valid/invalid) repeated-measures ANOVA yielded a main effect of Probability (F_(1.7,30.8)_ = 4.6, p = .023, η^2^ = .05, BF_incl_ = 5.4), reflecting overall slower responses in higher probability blocks. In terms of a main effect of Validity participants were slower for invalid compared to valid trials (F_(1,18)_ = 31.6, p < .001, η^2^ = .39, BF_incl_ > 100), and the interaction between Probability and Validity was also significant (F_(1.3,24.3)_ = 8.9, p = .003, η^2^ = .048, BF_incl_ = 7.9), reflecting that participants were slower in higher probability blocks in particular in invalid trials (see Figure 4). Thus, in our Pilot study, we were able to replicate the main finding by Vossel and colleagues (2014) in the context of internally generated predictions as well.

**Figure 4.**
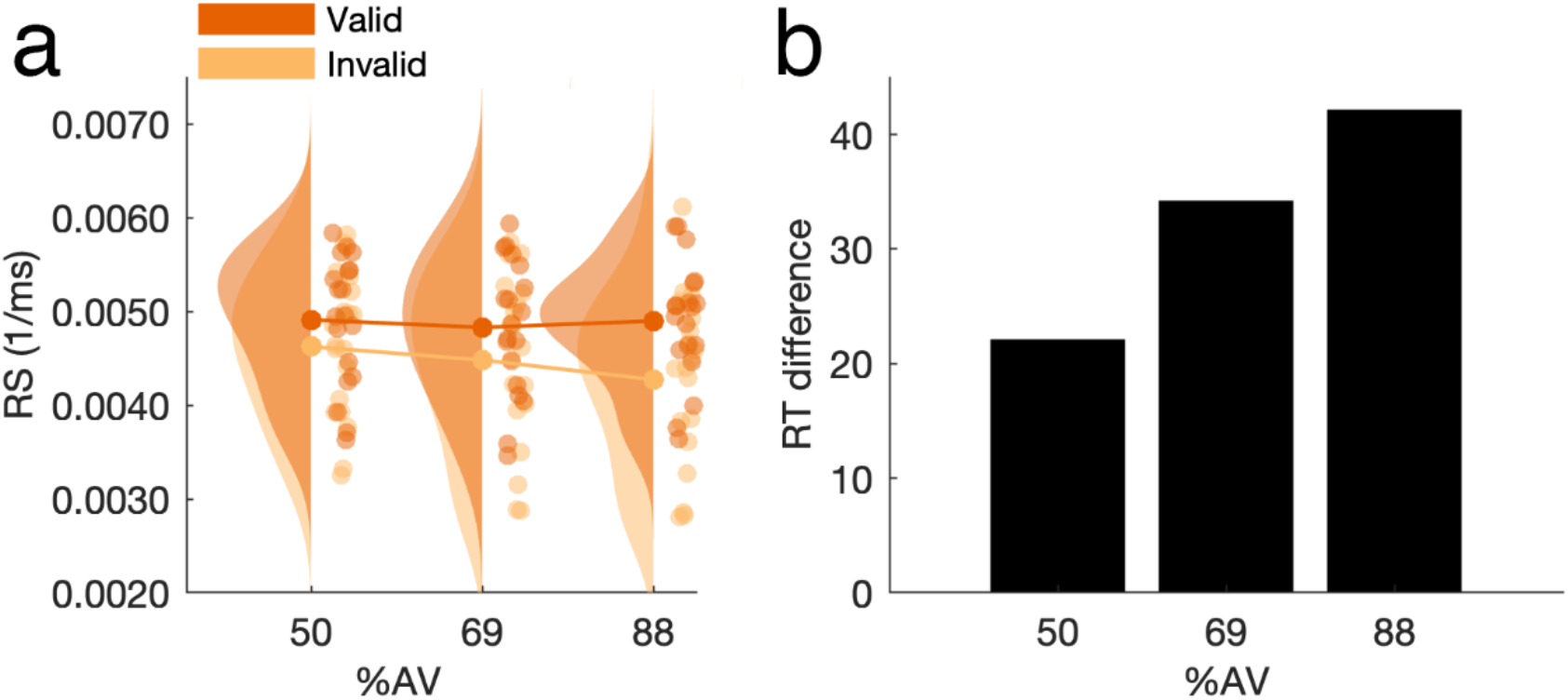
Eye movement results for the pilot study. Reaction speed across probability conditions split by valid and invalid trials (**a**), and the RT difference between valid and invalid trials in milliseconds (**b**). This figure shows that as the probability of a valid trial increased, the RT cost of a violated internally-generated expectation became larger.

#### HGF

In line with Vossel and colleagues (2014), we compared the three different response models by which the probability estimate for action validity could be transformed into RS using Bayesian model selection (Stephan et al., 2009). Rather than the precision response model, we found the belief response model to be most likely based on the protected exceedance probability (precision = .014, belief = .977, surprise = .01). The Bayes Omnibus Risk suggested all three models were not equally likely (p = .029). As can be seen in Figure 5, the trajectories for the averaged Bayesian parameters fit the presented validity evolution reasonably well. Thus, while for Vossel and colleagues (2014), the precision response model fitted best, in our case the optimal relationship between the allocation of attentional resources and predictability is a linear one. The question is whether this difference is specific to our action version of the experiment. To answer this question, we turn to our main experiment in which we directly contrast internally generated predictions through actions with externally induced predictions by cues.

**Figure 5.**
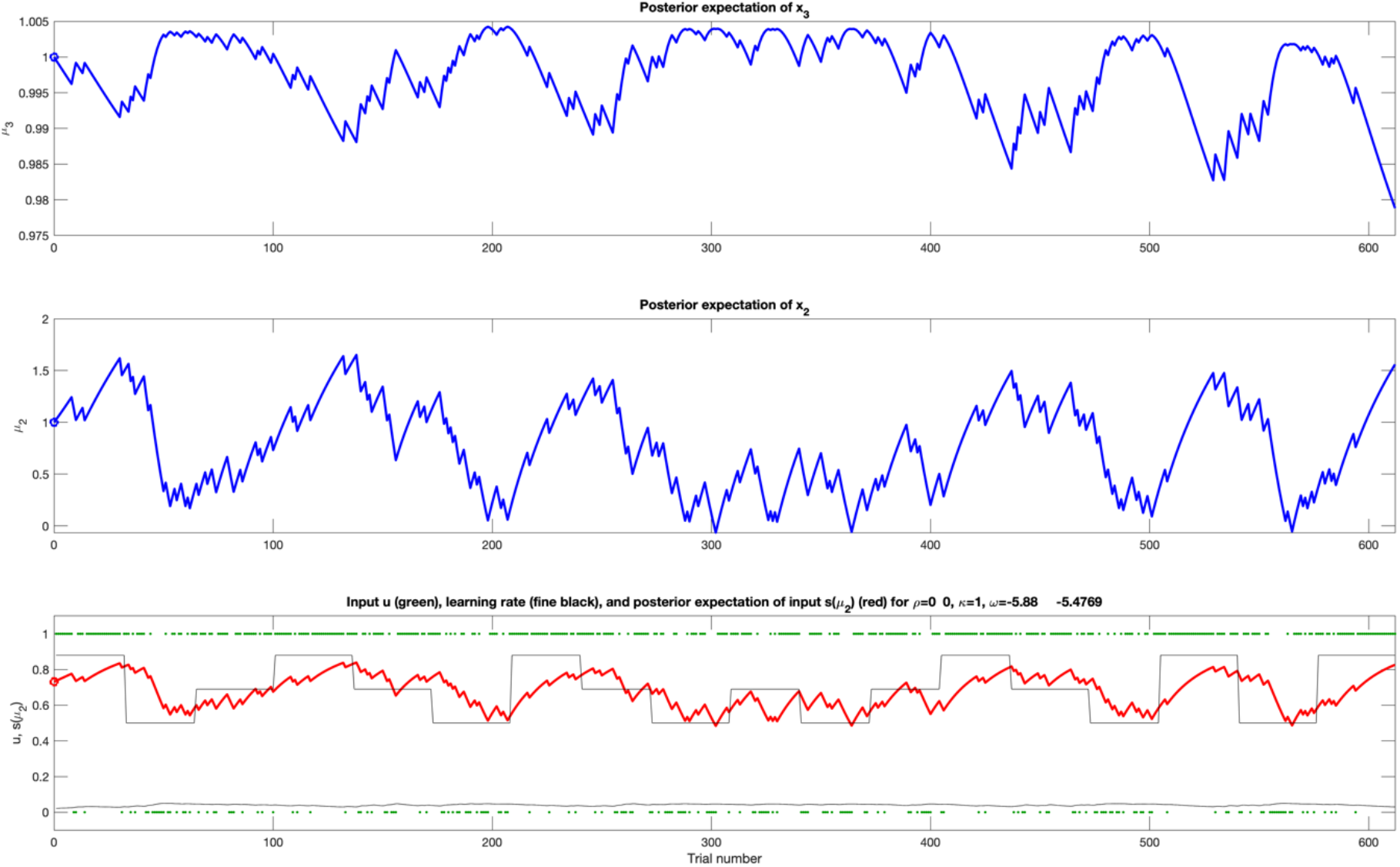
Trajectories for all three HGF levels derived from the averaged parameters belonging to the “belief” response model based on the Pilot data. The lowest panel shows the trial-by-trial probability estimate (red) against the actual probability blocks (black). It shows how the model successfully estimates and tracks the real probability relationship between action/cue and stimulus location.

### Main experiment

Similar to Vossel et al (2014), in total 21.6% of data was excluded from further analyses. 15.8% of trials were excluded because the saccade started more than 1° from fixation, 2.6% because saccadic RT was smaller than 90 ms, 2.1% due to less than two-thirds of the distance being traversed to one of either target locations, .53% because no saccade was detected, .26% because trials consisted of more than 20% missing values, and .16% due to incorrect trials.

#### Eye movement behavior

We first determined whether participants were overall faster on valid trials. The 3 (Probability; 88/69/50%) x 2 (Validity; valid/invalid) x 2 (Expectation; action-/cue-induced) repeated-measures ANOVA revealed a significant main effect of Validity (F_(1,24)_ = 59.3, p < .001, η^2^ = .15, BF_incl_ > 100), reflecting faster responses (higher RS) for valid compared to invalid trials. In addition, contrary to our preregistered prediction, participants became slower in higher probability blocks (F_(2,48)_ = 3.2, p = .048, η^2^ = .005, BF_incl_ = .08). A third main effect of Expectation indicated that participants were overall slower in the action compared to the cue condition (F_(1,24)_ = 6.3, p = .019, η^2^ = .14, BF_incl_ > 100).

We expected to find that expectations stemming from self-initiated actions would induce a larger reaction-time cost when violated compared to externally-cued expectations. Yet, the predicted significant three-way interaction between Probability, Validity, and Expectation was only trend level significant (F_(2,48)_ = 2.9, p = .066, η^2^ = .002, BF_incl_ = .15) (see Figure 6). We also did not find an interaction between Validity and Expectation (F_(1,24)_ = 2.7, p = .12, η^2^ = .002, BF_incl_ = .26). The same was true for the interaction between Probability and Validity, which was also trend-level significant (F_(2,48)_ = 2.6, p = .089, η^2^ = .002, BF_incl_ = .1). Thus, while we found a pattern of findings in the expected direction, it was not statistically robust.

**Figure 6.**
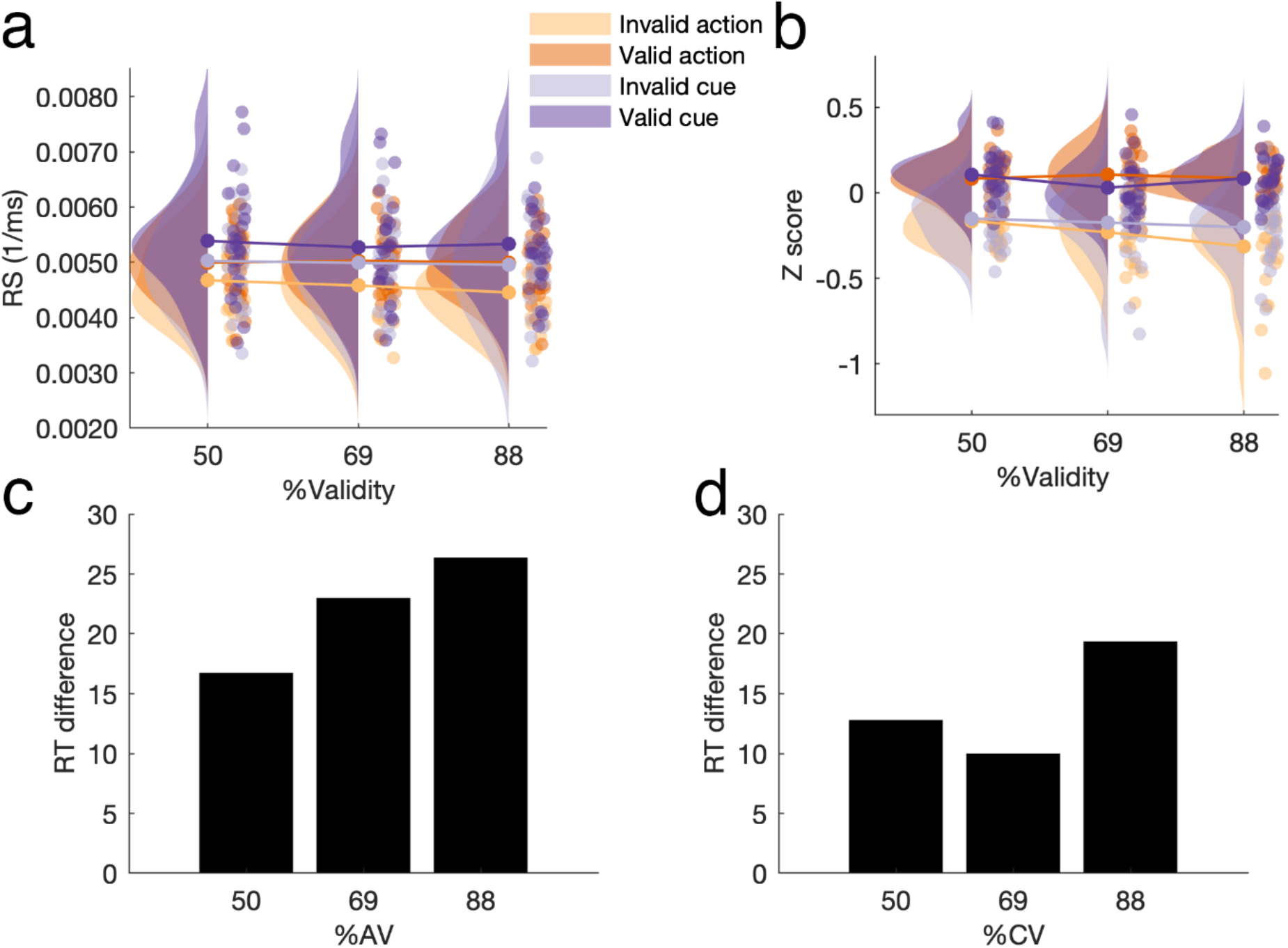
Eye movement results for the main experiment. Reaction speed across action/cue validity (AV/CV) conditions and split by **a**) valid and invalid trials, **b**) the same results but normalized to account for the overall difference in RS between the action and cue condition, and the reaction time difference between valid and invalid trials in milliseconds for the **c**) action and **d**) cue condition. While the RT cost for violated predictions is larger for 88% compared to 50% probability blocks in both conditions, we don’t find a consistent gradual increase in the cue condition as a function of cue validity.

#### Control analyses

We did not expect to find an effect of session order concerning whether the action or cue condition was completed first, but this expectation did not hold up. While we did not find a between-subject effect of Order (F_(1.23)_ = .34, p = .57, η^2^ = .009, BF_incl_ = .56), we did find a number of interactions. The interaction between Order and Expectation indicated that participants were faster in the cue session if they completed the action session first, but if the cue condition was completed first there was no difference (F_(2,23)_ = 4.7, p = .041, η^2^ = .034, BF_incl_ > 100). Order interacted with Probability in that participants were slower in 88% probability blocks if they completed the action session first, while they were fastest in 50% probability blocks when the cue session was completed first. (F_(2,46)_ = 6.5, p = .003, η^2^ = .003, BF_incl_ = .29). We also found a three-way interaction between Probability, Expectation, and Order (F_(2,46)_ = 3.7, p = .03, η^2^ = .002, BF_incl_ = .21), where participants were faster for all probabilities in the cue session only if they had already completed the action session. This difference between sessions across probabilities was absent if the cue session was completed first. Finally, we found a four-way interaction between Probability, Expectation, Validity, and Order (F_(2,46)_ = 3.3, p = .045, η^2^ > .001, BF_incl_ = .12), reflecting an enhanced three-way interaction between Probability, Expectation, and Validity if the action session was completed first. The task results outside Session Order reported above did not change in a meaningful way in this model. Given the relatively small sample size for this between-subject Order analysis, and the unexpected direction of some of the Order effects observed, it is unclear how to weigh these results.

In line with Vossel and colleagues (2014), but in addition to our preregistration, we conducted a control analysis for Time, in which we added a within-subject factor contrasting the first and second half of each session. This analysis yielded a main effect of Time (F_(1,24)_ = 102.6, p < .001, η^2^ = .25, BF_incl_ > 100), reflecting faster responses in the second half. Time interacted with Probability (F_(2,48)_ = 30, p < .001, η^2^ = .02, BF_incl_ = .2), to the point that participants were slower in 88% probability blocks in the first half compared to lower probability blocks, but they were faster in 88% probability blocks in the second half. Finally, we found a three-way interaction between Time, Probability, and Expectation (F_(2,48)_ = 5, p = .01, η^2^ = .004, BF_incl_ = .002), in that the difference in response time between the first and second half was larger for the action session, in particular for 50% and 88% probability blocks.

#### HGF

As to our Bayesian modelling results, we expected to replicate Vossel and colleagues (2014) in terms of the optimal response model (“precision”), in the cue as well as the action condition, although our pilot results identified the belief response model as the optimal response model for the action condition. For the action condition, Bayesian model selection indeed revealed that the precision response model was most likely (protected exceedance probability; precision = .34, belief = .32, surprise = .33), but the Bayes Omnibus Risk was sufficiently high (p = .91) to suggest that the three response models did not differ in a meaningful way. For the cue condition the belief response model came out on top (protected exceedance probability; precision = .29, belief = .48, surprise = .23), but in this case as well, the Bayes Omnibus Risk did not suggest a meaningful difference between models (p = .69).

We selected the parameters from the best model (precision for action condition, belief for cue condition) to establish whether belief updating differed under the influence of externally generated cues versus internally generated actions. Specifically, we compared the subjectspecific parameters ω and *ϑ* reflecting the speed of trial-wise belief updating concerning cue/action validity (ω) and the belief about the volatility of cue/action validity (*ϑ*). We expected participants to showcase a lower ω value in the cue condition, reflecting slower updating of predictions. Yet, contrary to our expectation a paired-samples t-test did not reveal a difference between the cue and action condition for ω_2_ (t_(24)_ = 1.1, p = .28, d = .003, BF_10_ = .03). However, we did find that participants believed volatility was higher in the action condition as reflected by ω_3_ (t_(24)_ = 2.7, p = .01, d = .3, BF_10_ = 3.9).

#### Explorative analysis

To account for the unexpected effects of (session) Order reported above, and to stick close to the original study by Vossel and colleagues (2014) in which participants only participated in one session of the cue condition, we conducted additional analyses in which we regarded the first session separately. Said differently, we regarded the first session as a between-subject study in which one group (n = 10) completed the cue session, while another group completed the action session (n = 15) to examine if we could replicate Vossel and colleagues (2014) with this more similar design.

##### Eye movement behavior

The 3 (Probability; 88/69/50%) x 2 (Validity; valid/invalid) x 2 (Expectation; action-/cue-induced) mixed ANOVA did reveal, as can be seen in Figure 7, the expected three-way interaction between Probability, Validity, and Expectation (F_(1.3,31)_ = 4, p = .044, η^2^ = .004, BF_incl_ = 5), reflecting that self-generated predictions through actions incurred a larger reaction time cost in particular in the high probability condition. Moreover, a trend-level significant between-subject main effect of Expectation was observed (F_(1,23)_ = 3.7, p =.068, η^2^ = .11, BF_incl_ = 68.9). Within-subject main effects were similar to the original analyses above: responses were slower in higher probability blocks (F_(1.5,35.2)_ = 5, p = .019, η^2^ = .007, BF_incl_ = 6.8) and on invalid trials (F_(1,23)_ = 46.7, p < .001, η^2^ = .066, BF_incl_ > 100). We furthermore found a significant interaction between Probability and Expectation (F_(1.5,35.2)_ = 7.3, p = .004, η^2^ = .001, BF_incl_ = 16.8), reflecting that only in the action condition, participants became slower in higher probability blocks. The interaction between Validity and Expectation was also significant (F_(1,23)_ = 5.4, p = .029, η^2^ = .008, BF_incl_ = 47.1), reflecting a larger reduction in response time for valid vs. invalid trials in the action condition. We did not find a significant interaction between Probability and Validity (F_(1.3,31)_ = 1.7, p = .2, η^2^ = .002, BF_incl_ = 1.4). Thus, when we look solely at the first session, we find that violation of predictions through invalid trials incurred a larger reaction time cost when a prediction stems from action rather than from a sensory cue, in particular under high probability conditions.

**Figure 7.**
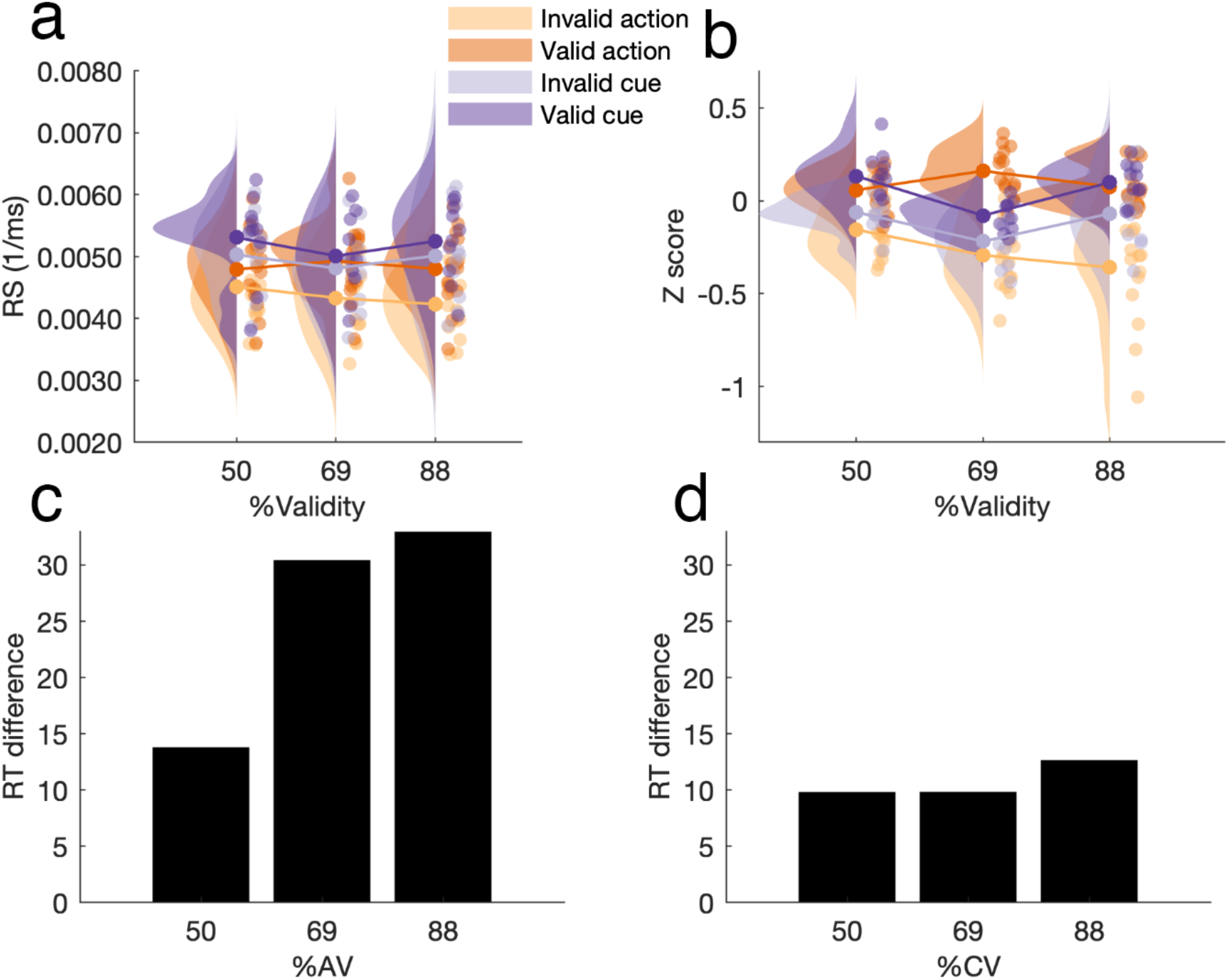
Eye movement results for only the first session of the main experiment. Reaction speed **a**) across probability conditions and split by valid and invalid trials, **b**) the same results but normalized, and the reaction time difference between valid and invalid trials in milliseconds for the **c**) action and **d**) cue condition. The RT cost for violated predictions is larger for 88% compared to 50% probability blocks in the action condition, but this difference is practically absent in the cue condition.

##### HGF

We repeated the procedure to select the best response model for the first session only. For the action condition, the surprise response model was most likely (protected exceedance probability; precision = .32, belief = .32, surprise = .35), but there was again little indication to assume the difference between these models was meaningful as reflected by the Bayes Omnibus Risk (p = .87). For the cue condition Bayesian model selection suggested the belief model was again best (protected exceedance probability; precision = .36, belief = .41, surprise = .23), but the Bayes Omnibus risk here too did not suggest a meaningful difference (p = .68)

We once again took the subject-specific parameters ω and *ϑ* from the winning models to compare them between the cue and action condition. An independent samples t-test did not reveal a difference for either ω_2_ (t_(23)_ = .83, p = .42, d = .34, BF_10_ = .48) or ω_3_ (t_(23)_ = .25, p = .8, d = .1, BF_10_ = .38). Thus, when controlling for time by reducing our analyses to only the Session 1 data, our modelling results did not change in a meaningful way.

##### Controlling for time

For the first session only as well, and in line with Vossel and colleagues (2014), we controlled for differences between the first and second half of the experiment by including a within-subject factor Time in our ANOVA. This resulted in a 2 (Time; first/second half) x 3 (Probability; 88/69/50%) x 2 (Validity; valid/invalid) x 2 (Expectation; action-/cue-induced) mixed ANOVA. Participants were again significantly faster in the second half of the session compared to the first half (F_(1,23)_ = 82.4, p < .001, η^2^ = .2, BF_incl_ > 100). Time interacted with Probability (F_(1.8,43.1)_ = 19.2, p < .001, η^2^ = .02, BF_incl_ > 100): responses became slower with higher probability blocks, but only in the first half of the session. Time did not interact with other variables (all p > .12). Said differently, unlike Vossel and colleagues, we still found differences between the first and second half of the experiment, but these did not differ between the action and cue groups.

## Discussion

In the present study, we addressed the following question: how does the response to induced predictions and their violation differ when these predictions stem from externally induced sensory cues compared to internally generated actions? We hypothesized that self-generated predictions would incur a larger reaction time cost when violated compared to cue-induced predictions, given that self-generated predictions allow a system to engage actively in testing and revising its models instead of being at the whim of the external environment to do so. We explored this question specifically within the context of predictive processing, and under the assumption that our participants perform approximate Bayesian inference. In absolute terms we found supporting evidence for our hypothesis: participants showcased a larger reaction time difference between valid and invalid trials in the action condition, particularly for high-probability trials, but it must be noted that the main 3-way interaction we were after remained at trend level significance for our main preregistered analyses, and only became statistically significant once we exploratively isolated the first session to account for unexpectedly observed session order effects.

In terms of modelling, only in our pilot data were we able to establish a difference between response models through a significant Bayes Omnibus Risk, but even then, it was not the “precision” model that was optimal, as we expected to find based on the study by Vossel and colleagues (2014). In terms of model parameters, the difference between action and cue conditions was reflected in the volatility parameter ω_3_ in our main analyses instead of the hypothesized speed of belief-updating parameter ω_2_. This difference disappeared for our explorative analysis isolating only the first session, possibly due to the small groups being compared (N = 10 and N = 15). Overall, if these behavioral and modelling results are to be believed, it does seem that the larger reaction time cost induced by the violation of action-induced predictions translates to participants becoming more sensitive to the volatility of relevant probability relationships in their surroundings.

A difference between how stimuli stemming from internally generated and externally induced predictions affect model updating was expected based on the idea that self-generated predictions are generally afforded more weight compared to externally induced predictions (Brown et al., 2013), because they allow a system to put its own models to the test, instead of being reliant on external change in its environment. This idea aligns with the way fine-tuning of behavior is understood within the context of sensorimotor learning. As in the case of male songbirds who in a first “sensory” stage store a song-template in memory, only to put this template to the test in a second “sensorimotor” stage where they actively but gradually approach the stored template (Brainard & Doupe, 2000). In our experiment, the “template” is the intuitive relationship between a left and right action with a left and right visual effect which is put to the test repeatedly. A problem with transposing this example to the visual domain is that it is unclear whether there are anatomical connections directly connecting the motor cortex to the visual cortex, as is the case for the auditory cortex (Reznik & Mukamel, 2019). Nevertheless, it could be that such a process takes place between the motor cortex and more abstract representations such as “left” and “right” instead of purely visual representations per se. Indeed, such a proposal would fit with predictive processing accounts of sensory attenuation where neural responses to predicted stimuli are reduced for aggregated signals across many neurons, possibly reflecting a sharpening of the relevant neural representation (Bell et al., 2016; Kok et al., 2012; Press et al., 2020; Reznik & Mukamel, 2019), especially when predictions are self-generated (Brown et al., 2013; Korka et al., 2019, 2022). It is possible that increased sensitivity to violation for self-generated predictions comes from increased sharpening given that self-generated predictions are weighted more heavily by the system in question.

Several limitations pertain to the present study. Unlike Vossel and colleagues (2014), we found an effect of Time we did not hypothesize: participants were faster in the second half of the experiment compared to the first. One difference between our and the original study is that ours took longer to complete. Whereas Vossel and colleagues report their experiment lasting 35 minutes (p. 1442), ours took over an hour on average to complete. In the original study, the authors report allowing one and four short rest periods across their two datasets, which were increased from the first to the second to increase data quality. With this in mind, we allowed for eight short rest periods, which will have lengthened our experiment. More importantly however, participants in the study by Vossel and colleagues (2014) responded faster and made more errors (~5% vs < 1%). This difference may indicate a difference in speed-accuracy tradeoff between the studies. As such, future research should emphasize speed over accuracy in attempted replications like these. More practice trials may reduce the time effect as well by eliminating task familiarization during experimental trials.

We also found a session order effect we did not hypothesize. Participants’ performance was faster in the cue session after completing the action session first, while participants were not faster in the action session after having completed the cue session first. Apparently, a task like this is learned more readily in the cue condition. This makes intuitive sense because the cue condition required less from participants. In the cue session, participants merely had to sit still and move their eyes to the target when it appeared, whereas in the action session they also had to choose an action every single trial. We chose a within-subject design to exploit the increase in statistical power and reduction in random noise these designs are privy to, but given these order effects, it may be better to opt for a between-subject design in future research. Care should be taken in recruiting enough participants given that even with our within-subject design, our effects of interest remained at trend level.

In addition to the limitations that pertain to our basic behavioral results, we were unable to replicate the modelling results by Vossel and colleagues (2014). The difference resides in the best fitting response model, which stands for the way the trial-by-trial probability of cue or action validity is transposed to response speed through the computation of the attentional factor α. In the original study this transposition was best governed by precision (π^(t)^) in line with the proposal by Feldman and Friston (2010), but in our case only for our pilot data did we find a clear winning model (“belief”). While the behavioral limitations discussed above may have contributed to these ambiguous modelling results, there are additional angles to consider. Whereas extensive model comparison was performed in the original study (Vossel, Mathys, et al., 2014), in subsequent studies employing the same task an optimal “precision” response model is assumed without repeated model comparisons for these new datasets (Vossel, Bauer, et al., 2014; Vossel et al., 2015), while in other work with different attention tasks, it was not the “precision” but the “belief” model that fitted best (Dombert et al., 2016; Kuhns et al., 2017). It’s possible that the tasks used were sufficiently different to warrant a different optimal response model, but it is also possible that the model fitting procedure is not as robust as we may desire. The Vossel et al. (2014) study had an even smaller sample size than the current study, which may have led to less robust results. Future research would do well to elucidate under what conditions different response models fit best with larger samples sizes. Perhaps a better modelling approach altogether to test the current research question would be one in which the difference is considered between perceptual and active inference, as they’ve been theorized to optimize two separate free energy functionals (Parr & Friston, 2019).

While the present study leaves room for improvement in experimental design, the results we report nevertheless uphold the importance of considering the difference between prediction of self-generated and externally imposed stimuli, especially within the context of predictive processing. While both perception and action serve the same goal: the minimization of prediction error or free energy (Clark, 2013; Friston, 2010; Hohwy, 2013), it seems likely that self-generated predictions are afforded more weight by the generating system. It may be worth considering letting go of some experimental control for the sake of an inside-out perspective.

## Acknowledgements

We would like to thank Simone Vossel for helpful modelling guidance.

